# The causal structure of age-dependent limbic decline: fornix white matter glia damage causes hippocampal grey matter damage, not *vice versa*

**DOI:** 10.1101/440917

**Authors:** Claudia Metzler-Baddeley, Jilu P. Mole, Rebecca Sims, Fabrizio Fasano, John Evans, Derek K. Jones, John P. Aggleton, Roland J. Baddeley

## Abstract

Aging leads to gray and white matter decline but their causation remains unclear. We explored two broad classes of models of age and dementia risk related brain changes. The first class of models emphasises the importance of gray matter: age and risk-related processes cause neurodegeneration and this causes damage in associated white matter tracts. The second class of models reverses the direction of causation: aging and risk factors cause white matter damage and this leads to gray matter damage. We compared these models with linear mediation analysis and quantitative multi-modal MRI indices (from diffusion, quantitative magnetization transfer and relaxometry imaging) of tissue properties in two limbic structures implicated in age-related memory decline: the hippocampus and the fornix in 166 asymptomatic individuals (aged 38 - 71 years). Aging was associated with apparent glia but not axon density damage in the fornix. Mediation analysis unambiguously supported white matter damage causing gray matter decline; controlling for fornix glia damage, the correlation between age and hippocampal damage disappears, but not *vice versa*. Fornix and hippocampal tissue loss were both associated with reductions in episodic memory performance. The implications of these findings for neuroglia and neurodegenerative models of aging and late onset dementia are discussed.

## Introduction

The world’s population is growing older and an increasing number of people over the age of 65 years will develop cognitive impairment due to late onset Alzheimer’s disease (LOAD)^1^. The pathological processes leading to LOAD accumulate over many years and share some overlapping features with normal aging processes^2^. For example, amyloid-β plaques and neurofibrillary tangles also occur in brains of non-demented older people^3–5^. Recent recommendations by the Lancet commission highlight the importance of dementia prevention studies that start in midlife with the aim of developing effective therapeutics in the future^6^. To achieve this, however, it is necessary to identify those individuals that will develop LOAD prior to the onset of memory impairment. This in turn requires a better understanding of how aging and risk factors of LOAD affect the brain in cognitively healthy individuals from midlife onwards.

Aging causes damage to both gray and white matter in the human brain. White matter consists of myelinated and unmyelinated axons and neuroglia cells, i.e., oligodendrocytes, astrocytes and microglia, while gray matter comprises neuronal cell bodies, synapses, dendrites and glia. Neuroglia cells are essential for normal synaptic and neuronal activity as they maintain the brain’s homeostasis, form myelin, protect neurons, and are dynamically remodelled to support brain plasticity ^7–10^. There are two classes of causal models of aging and LOAD risk: the first class of neurodegenerative models proposes that neuronal and synaptic loss precede white matter glia and axon damage. According to an influential neurodegenerative model, the amyloid cascade hypothesis ^11, 12^, aging and LOAD risk factors lead to metabolic changes in the amyloid precursor protein that result in the aggregation of amyloid-β plaques, which trigger pathological events including the formation of neurofibrillary tangles, loss of synapses, neurons and their axons. Neuroglia changes such as reactive microglia occur in response to increased plaque and tangle burden ^13–15^. The second class of model reverses the direction of causality, stating that the normal aging process in interaction with genetic and lifestyle risk factors of LOAD will cause neuroglia damage, resulting in impaired myelination, reduced microglia-mediated clearance and neuroinflammation, which in turn instigates pathological processes that lead to abnormal metabolism in key proteins and neuronal death ^16–19^. The two classes of models predict opposite patterns of age and risk-related changes in white and gray matter. While in neurodegenerative models, white matter damage follows gray matter neuronal loss, the neuroglia model predicts that white matter neuroglia damage causes gray matter tissue loss.

**Table 1:**
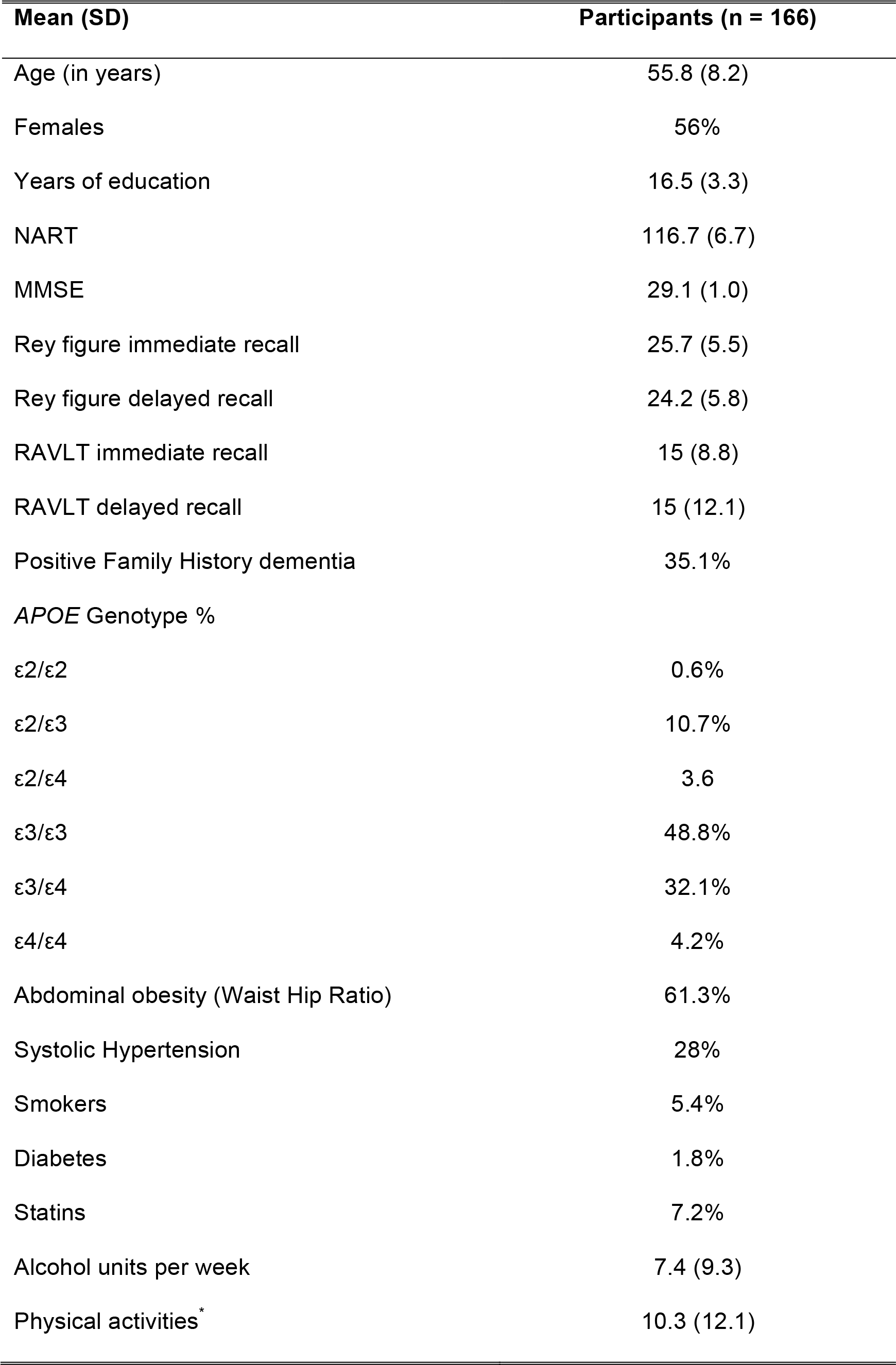
Summary of demographic, genetic and lifestyle risk information of participants.

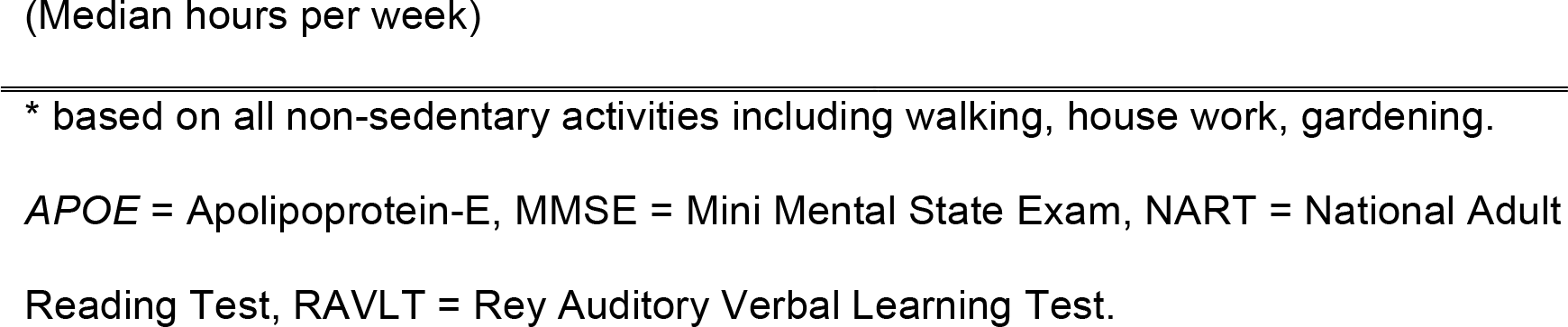

**Figure 1.**
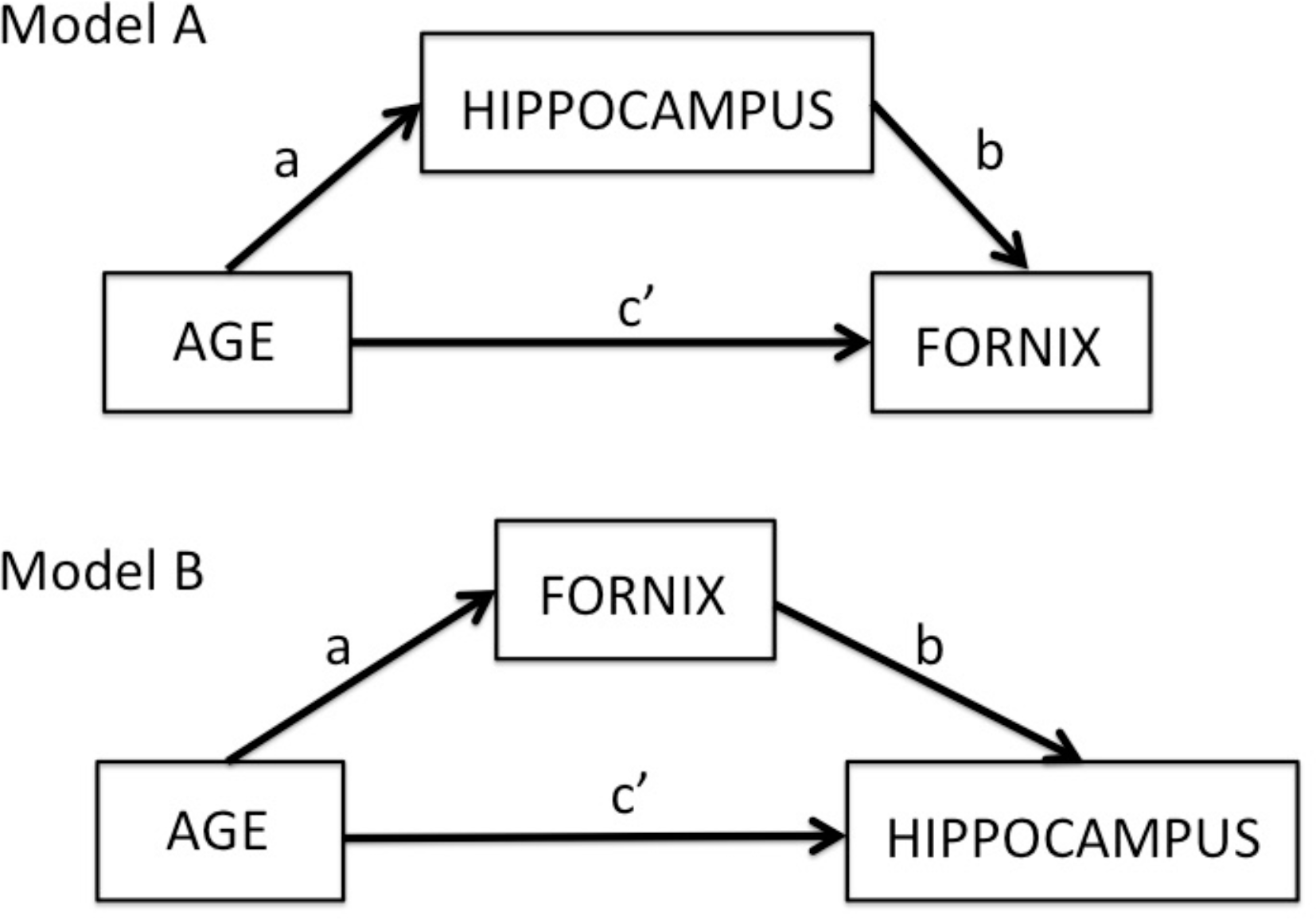
Overview of two mediation models: Model A tests for the indirect effects of hippocampal mediator variables on the direct effect of age on the fornix (path c’). The indirect effect is the product a*b of the correlations between age and hippocampus (path a) and the partial correlation between hippocampus and fornix accounting for age (path b). Model B tests the indirect effects of the fornix mediator variables on the direct effect of age on the hippocampus.

Currently, the nature of age-related white and gray matter tissue changes and the causal direction between them remain unknown. This study therefore investigated the impact of age and risk on novel quantitative MRI indices of apparent neurite and glia properties, and discriminated between the two types of models using linear mediation analysis^20–22^. More specifically, we investigated the effects of aging and risk factors on tissue properties of two important gray and white matter structures involved in age-related memory decline, the hippocampus^23^ and the fornix^24^, in 166 cognitive healthy individuals (39-71 years old) (Table1). Linear mediation analysis is the simplest example of a rapidly developing methodology to study, not only the correlations between variables, but also the causal structure between them ^25^.

Assuming that age is correlated with both hippocampal and fornix metrics and that these in turn are correlated with each other, it is possible with linear regression to separate any direct effects of, for example, age on the fornix (Figure 1 path c’) from any indirect effects (Figure 1 path a*b) caused by the effect of age on the hippocampus (Figure 1 path a) and the effect of the hippocampus on the fornix when age is controlled for (Figure 1 path b). In other words, it is possible to assess a) whether age-related hippocampal differences mediate the relationship between age and fornix tissue properties (Model A in Figure 1) and b) whether age-related fornix differences mediate age effects on hippocampal tissue properties (Model B in Figure 1). In most situations in the social and medical sciences, the results of a mediation analysis are an estimate of the relative effect sizes of the direct and the indirect mediating relationships. However, in rare circumstances a full mediation occurs, i.e. controlling for the effects of mediation results in the removal of the correlation due to the direct pathway c’. When full mediation is only observed in one direction, it is possible to infer about the causal chain of the correlations between the specific variables under investigation. For example, if full mediation is observed in Model A (Figure 1) such that the inclusion of hippocampal mediators removes the age effects on the fornix but is not present in Model B, such that fornix mediators do not fully mediate age effects on hippocampal properties (Figure 1), then this would indicate that hippocampal gray matter differences were causing white matter fornix differences. This pattern of results would be consistent with neurodegenerative models of aging. However, if full mediation occurs in Model B but not Model A, then this would indicate that age-related fornix differences were causing hippocampal gray matter differences. This pattern of results is consistent with neuroglia models of aging.

**Figure 2.**
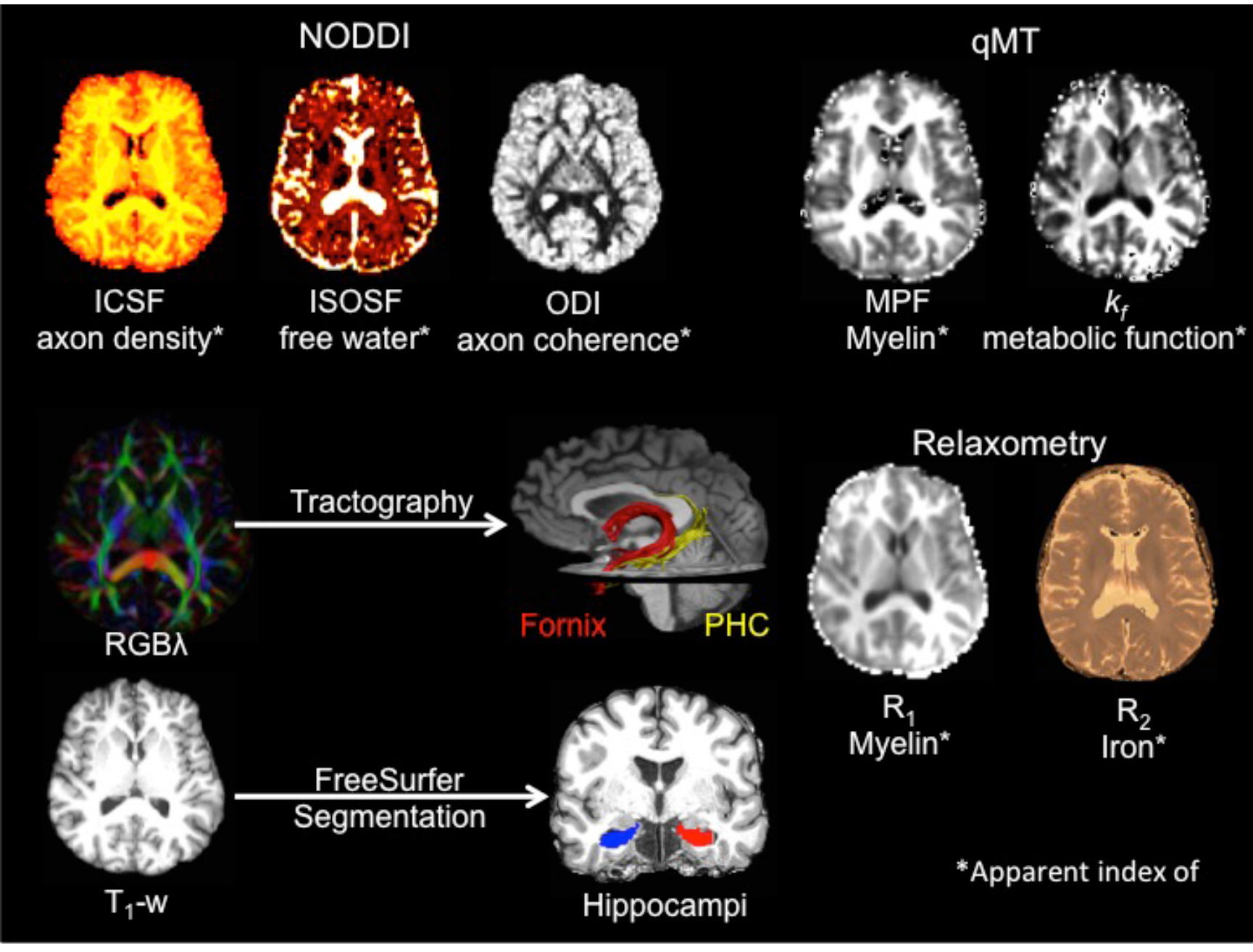
Displays the MRI modalities and maps acquired from dual-shell high angular resolution imaging (HARDI), quantitative magnetisation transfer (qMT) imaging and from T_1_ and T_2_ relaxometry. HARDI data were modelled with neurite orientation dispersion and density (NODDI) yielding maps of intracellular signal fraction (ICSF), isotropic signal fraction (ISOSF) and orientation density index (ODI). qMT based maps included the macromolecular proton fraction (MPF) and the forward exchange rate *k*_f_. Maps from relaxometry were the longitudinal relaxation rate R_1_ (1/T_1_) and R_2_ (1/T_2_). Mean indices of the metrics were extracted from left and right hippocampi, fornix and parahippocampal cinguli (PHC) tracts. Hippocampi were segmented from T_1_-weighted images with FreeSurfer version 5.3 and fornix and PHC were reconstructed with damped-Richardson Lucy spherical deconvolution (dRL) based deterministic tractography on colour coded principal direction maps (RGB*λ*).

Quantitative magnetic resonance imaging (MRI) can provide *in vivo* indices of biophysical white and gray matter tissue properties. We used dual-shell high angular resolution diffusion (HARDI) ^26^, quantitative magnetization transfer (qMT) ^27–30^ and T_1_ and T_2_-relaxometry MRI to assess tissue properties in the hippocampus, the fornix and the parahippocampal cingulum (PHC), as a temporal lobe comparison pathway. Separate estimates of axon and glia-related microstructural properties of white matter were obtained from the neurite orientation dispersion and density imaging (NODDI) ^31^ and the qMT ^32–35^ models (Figure 2). NODDI was fitted to the HARDI data to separate different compartments of the diffusion signal. Restricted anisotropic non-Gaussian diffusion in the intra-axonal space was quantified with the intracellular signal fraction (ICSF), an index of apparent axon density. Free isotropic Gaussian diffusion was estimated with the isotropic signal fraction (ISOSF), which provides an estimate of cerebrospinal fluid (CSF) based partial volume contribution to the diffusion signal. In addition, NODDI models the orientational coherence of white matter fibers with a Watson distribution yielding the orientation dispersion index (ODI). Diffusion MRI based indices are well-known measures of white matter microstructure ^36–38^ and NODDI ICSF and ODI have been proposed to be particularly sensitive to axonal density and dispersion ^39, 40^.

These axon-related microstructural indices from NODDI were complemented with qMT metrics that provide improved white matter glia specificity compared to diffusion weighted MRI ^32–35^. In qMT, free water and semisolid macromolecular constituents of tissue are estimated by applying off-resonance radiofrequency pulses with constant amplitudes that are varied between offsets, to selectively saturate the macromolecular magnetization with exchange processes resulting in magnetization transfer between saturated macromolecules and free water ^32^. In white matter, magnetization transfer is dominated by myelin^41, 42^, and is also sensitive to microglia-mediated inflammation ^33, 43, 44^. Here we utilised i. the relative number of spins in the macromolecular pool, the macromolecular proton fraction (MPF) as an index of white matter neuroglia that will largely reflect its myelin content and ii. the forward exchange rate *k*_*f*_, an index of the rate of the magnetization transfer process ^32^, that has been proposed to reflect metabolic efficiency of mitochondrial function ^45^ (Figure 2). Finally, the longitudinal relaxation rates R_1_ (1/T_1_) and R_2_ (1/T_2_) were acquired as additional estimates of the relative contribution of water, myelin and iron content of gray and white matter tissue ^46–49^ (Figure 2).

Mean values of all MRI indices were extracted for the white matter in the fornix and the left and right PHC. Tracts were reconstructed with spherical deconvolution-based deterministic tractography in ExploreDTI (version 4.8.3) ^50^ (Figure 2). Mean values of all metrics, except ICSF and ODI, which so far have only been validated in white matter ^31^, were extracted bilaterally from FreeSurfer (version 5.3) segmentations of the whole hippocampi ^51, 52^ These hippocampal regions include areas of the presubiculum, subiculum, cornu ammonis subfields 1-4, dentate gyrus, hippocampal tail and fissure but exclude cortical regions such as the entorhinal cortex ^53, 54^ (Figure 2).

To study how age-related differences in above MRI metrics relate to genetic and lifestyle risk factors of LOAD as well as to individual differences in episodic memory performance, we also acquired the following information (Table 1): Genetic risk was assessed by carriage of the Apolipoprotein-E (*APOE*) ε4 allele^55, 56^ and separately by a positive family history of a first grade relative with dementia of the Alzheimer’s, Lewy body or vascular type. In particular, we predicted more pronounced white matter glia reductions in *APOE* ε4 homozygotes (ε4/ε4) and heterozygotes (ε3/ ε4) than in ε2 and ε3 homozygotes (ε2/ε2, ε3/ε3) and heterozygotes (ε2/ε3), as cholesterol transport for myelin repair in the brain has been proposed to be less efficient in ε4 carriers ^56–59^ (ε2/ε4 carriers were excluded from the analysis). In addition, we assessed lifestyle related risk factors associated with the metabolic syndrome, notably central obesity, hypertension, alcohol consumption and physical inactivity ^60, 61^ (Table 1). We were particularly interested in the effects of central adiposity as obesity: i. is globally on the rise; ii. is known to be associated with chronic inflammation, insulin resistance and vascular problems ^60, 61^; iii. has been shown to accelerate brain aging of white matter ^62^; and iv. is associated with impaired fornix microstructure ^63^. Finally, episodic memory abilities were assessed with standard neuropsychological tests of verbal and non-verbal recall^64, 65^ (Table 1).

The neuroglia model predicts that aging and risk factors will primarily reduce glia sensitive metrics of MPF, *k*_*f*_, R_1_ and R_2_ whilst white matter microstructural indices of apparent axon density (ICSF) and orientational dispersion (ODI) should not be affected. Age and risk-related increases in ISOSF, reflecting increased CSF partial volume due to unspecific tissue loss, are predicted by both models. Importantly though, the neuroglia model predicts that age-related differences in glia sensitive metrics of the fornix will mediate age-related hippocampal differences (Model B), whilst the neurodegenerative model predicts that hippocampal differences will mediate age-related differences in the fornix (Model A).

## Results

### Omnibus multivariate regression analysis

Multivariate regression analysis with all hippocampal, fornix and PHC MRI outcome metrics as dependent variables (Figure 2), were tested simultaneously for omnibus effects of the following independent variables:

- age,
- genetic risk: family history of dementia (yes/no), *APOE* genotype [ε2 (ε2/ε2, ε2/ε3), ε3(ε3/ε3), ε4 (ε3/ε4, ε4/ε4)],
- lifestyle-related risk: central obesity assessed with the waist hip ratio (WHR), systolic and diastolic blood pressure, weekly alcohol consumption, physical activity and,
- potentially confounding variables of sex, years of education, and head size assessed with intracranial volume (ICV).

This analysis revealed significant omnibus main effects for age [F(31, 98) = 3.1, p = 0.000013; ηp^2^ = 0.49], ICV [F(31, 98) = 2.2, p = 0.002, ηp^2^ = 0.41] and sex [F(31, 98) = 1.7, p = 0.027, ηp^2^ = 0.35], but no other significant effects.

A *post-hoc* multivariate covariance analysis was then carried out to test for the effects of age on each of the hippocampal, fornix and PHC MRI indices separately whilst controlling for ICV and sex. All multiple comparisons were corrected with a False Discovery Rate (FDR) of 5%.

There were significant effects of age on:

- fornix MPF [F(2, 149) = 12.8, p = 0.000008, ηp^2^ = 0.15],
- fornix R_1_ [F(2, 149) = 12.3, p = 0.00001, ηp^2^ = 0.14],
- fornix *k*_*f*_ [F(2, 149) = 10.5, p = 0.00006, ηp^2^ = 0.12],
- fornix ISOSF [F(2, 149) = 8.55, p = 0.0003, ηp^2^ = 0.1],
- fornix ODI [F(2, 149) = 4.3, p = 0.015, ηp^2^ = 0.06],
- left PHC R_2_ [F(2, 149) = 6.5, p = 0.002, ηp^2^ = 0.08],
- right PHC R_1_ [F(2, 149) = 5.2, p = 0.007, ηp^2^ = 0.07],
- hippocampal ISOSF [left: F(2, 149) = 9.5, p = 0.00012, ηp^2^ = 0.11; right: F(2, 149) = 5.8, p = 0.004, ηp^2^ = 0.07],
- left hippocampal R_1_ [F(2, 149) = 5.4, p = 0.006, ηp^2^ = 0.07]
- left hippocampal *k*_*f*_ [F(2, 149) = 6.4, p = 0.002, ηp^2^ = 0.08]. (Figure 4).

No age effect was present for fornix ICSF [F(2, 149) = 2, p = 0.14, ηp^2^ = 0.026].

**Figure 3.**
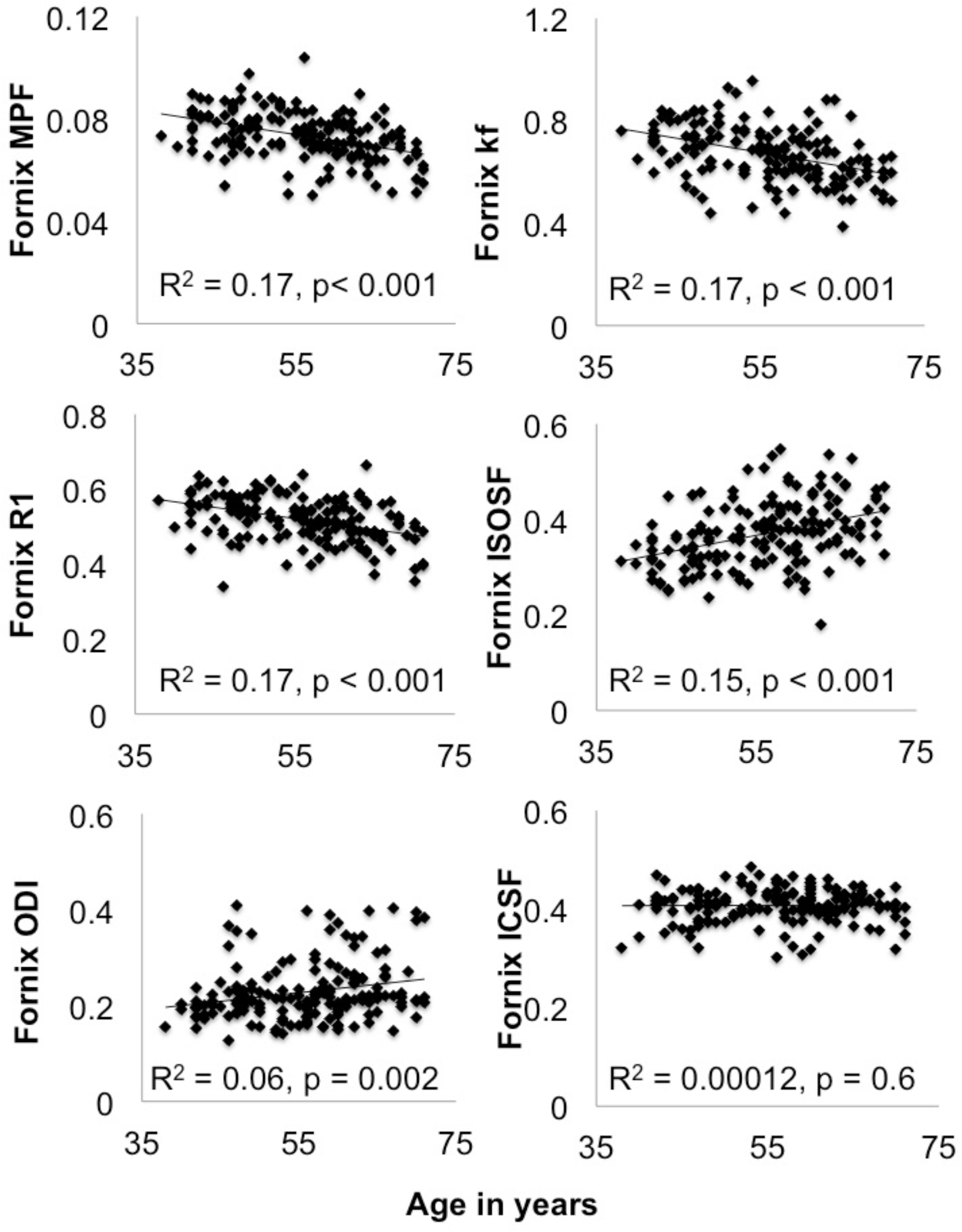
Plots the 5% false discovery rate corrected significant correlations between age and fornix macromolecular proton fraction (MPF), longitudinal relaxation rate R_1_, forward exchange rate *k*_*f*_, isotropic signal fraction (ISOSF) and orientation dispersion index (ODI) as well as the null-correlations with intracellular signal fraction (ICSF) for comparison.

**Figure 4.**
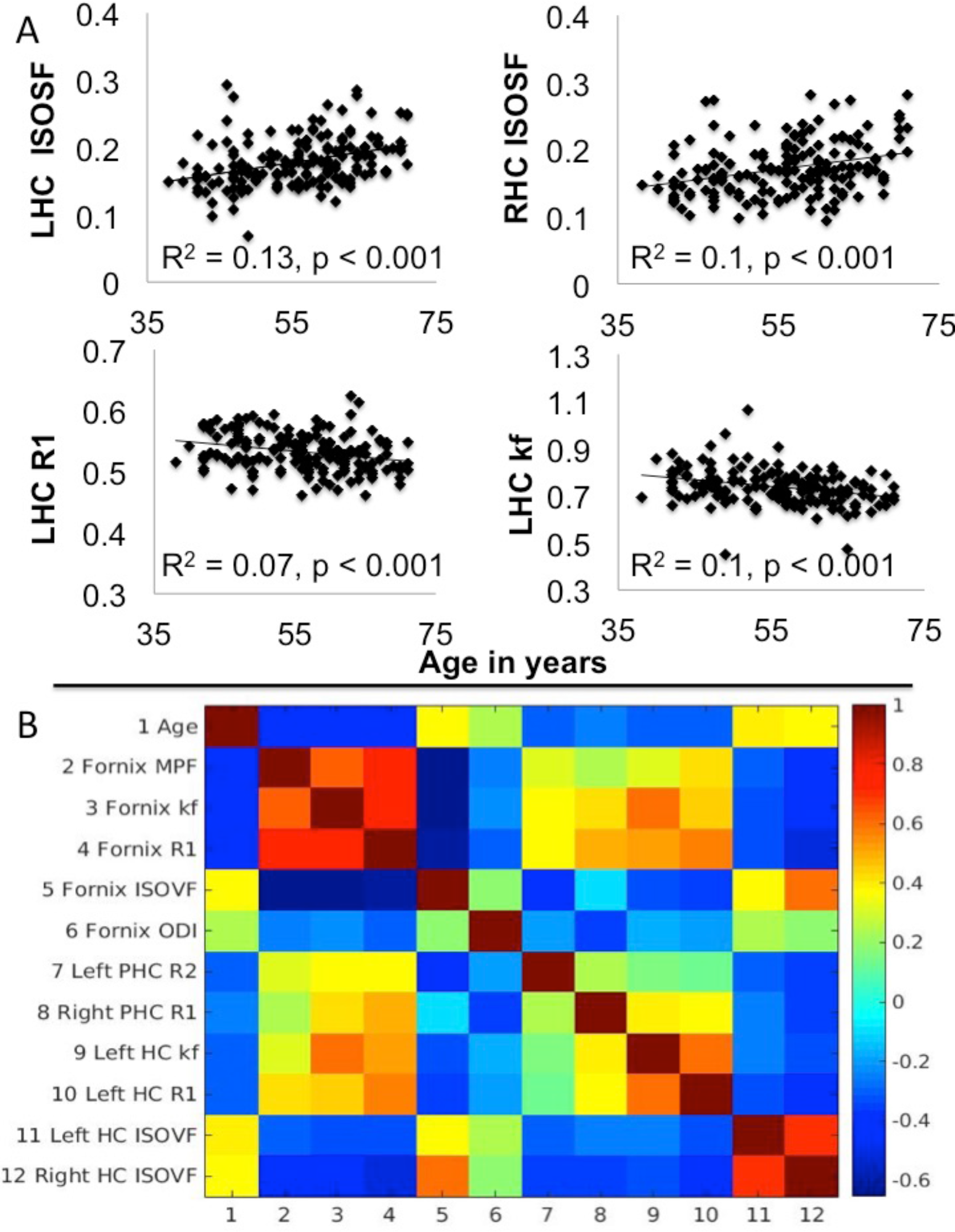
*A)* plots the 5% false discovery rate corrected significant correlations between age and left (L) and right (R) hippocampal (HC) isotropic signal fraction (ISOSF), left hippocampal R_1_ and *k*_*f*_. B) displays the Spearman correlation coefficient matrix between age and those hippocampal, fornix and parahippocampal (PHC) MRI metrics that showed significant age effects.

To study the direction of these age effects as well as the correlations between white and gray matter metrics, partial Spearman rho correlations controlling for sex and ICV were calculated between age and the white matter and hippocampal metrics (Figures 3 and 4).

Significant negative correlations were present between age and:

- fornix MPF [rho (151) = −0.47, p = 0.0000000019],
- fornix R_1_ [rho (151) = −0.48, p = 0.0000000002],
- fornix *k*_*f*_ [rho (151) = −0.47, p = 0.0000000014],
- right PHC R_1_ [rho (151) = −0.22, p = 0.008],
- left PHC R_2_ [rho (151) = −0.36, p = 0.000006],
- left hippocampal *k*_*f*_ [rho (151) = −0.34, p = 0.000018]
- left hippocampal R_1_ [rho (151) = −0.32, p = 0.00005].

Positive correlations with age were observed with:

- fornix ISOSF [rho (151) = 0.40, p = 0.0000002],
- fornix ODI [rho (151) = 0.25, p = 0.002],
- hippocampal ISOSF [left: rho (151) = 0.46, p = 0.000000003; right: rho (151) = 0.39, p = 0.00000046].

However, age did not correlate with fornix ICSF [rho (151) = −0.04, p = 0.59] and this ‘null’ correlation differed significantly from all other age correlations with fornix metrics [z_min_ = 2.5, p_2-tailed_ = 0.01; z_max_ = 4.15, p_2-tailed_ < 0.001] (Figure 3).

Figure 4B displays the partial Spearman correlation coefficient matrix between age, fornix and PHC white and hippocampal gray matter metrics. As can be seen, white matter microstructural metrics correlated significantly with the hippocampal gray matter metrics in the expected directions. That means, that there were positive correlations between the same metrics measured in gray and white matter and between MPF, *k*_*f*_, R_1_ and R_2_ metrics respectively, but negative correlations between MPF, *k*_*f*_, R_1_ and R_2_ metrics on one hand and ISOSF and ODI metrics on the other.

**Figure 5.**
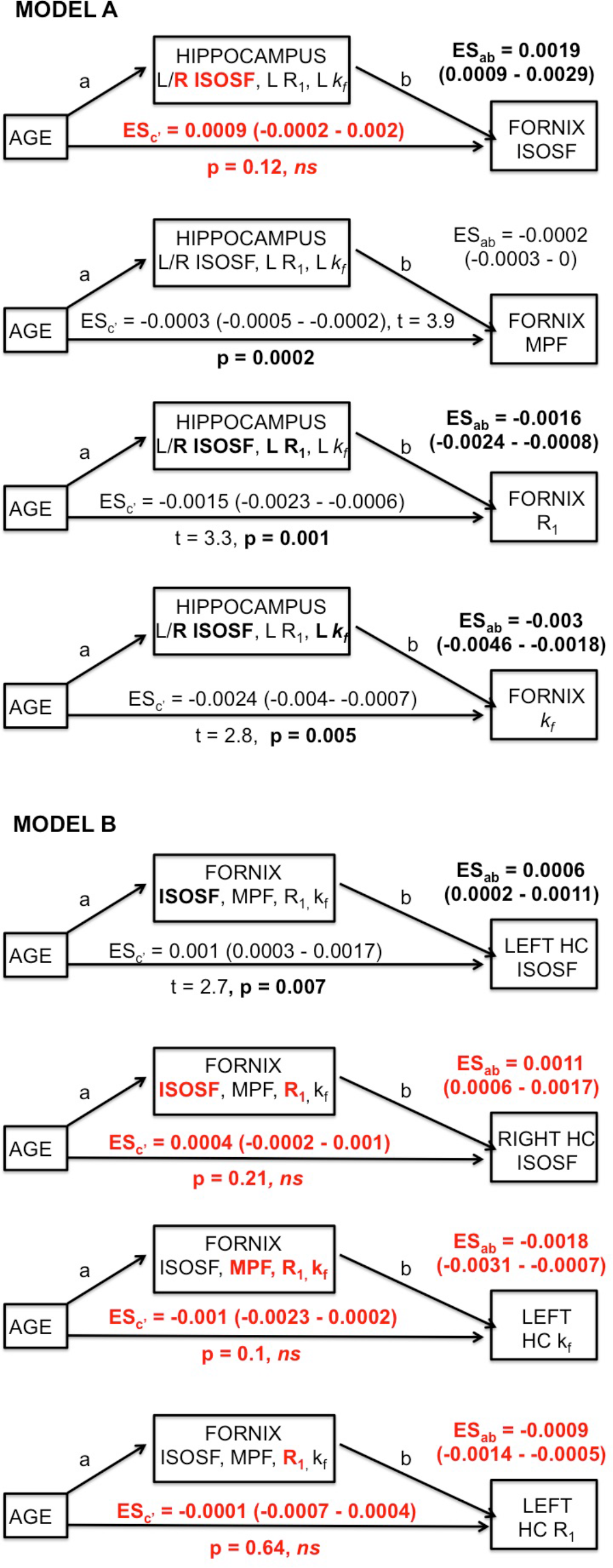
Summarises the results of the mediator analyses for Model A and B. 95% confidence intervals of the effect sizes (ES) were based on bootstrapping with 5000 replacements. Fornix mediators in Model B had significant indirect effects and fully mediated the direct age effects on right hippocampal isotropic signal fraction (ISOSF), left (L) hippocampal R_1_ and *k*_*f*_ (highlighted in red). In contrast, hippocampal mediators, although showing significant indirect effects (highlighted in bold), did not fully mediate the direct age effects on fornix MPF, R_1_ and *k*_*f*_. Right (R) hippocampal ISOSF fully mediated fornix ISOSF and *vice versa* but fornix mediators did not remove the age effect on left hippocampal ISOSF. Mediator variables that contributed significantly to the regression analyses after 5% false discovery rate correction are highlighted in bold.

### Mediation analyses between hippocampal and fornix metrics

To avoid any bias in the comparison of Model A and Model B, the number of mediator variables was kept constant. Model A included all hippocampal metrics that showed significant age effects, i.e., left and right hippocampal ISOSF, left hippocampal R_1_ and left hippocampal *k*_*f*_, as mediator variables for the direct correlations between age and fornix metrics. Model B included those four fornix metrics with the largest age effect sizes (as measured by rho) (ISOSF, MPF, *k*_*f*_, R_1_) as mediator variables for direct effects of aging on the hippocampus. There were no significant differences between the sizes of the age effects on fornix compared with hippocampal mediator variables (rho_max:age-fornix R1_ *versus* rho_min:age-left hippocampal R1_: z = −1.65, p_2-tailed_ = 0.099).

Figures 5A and 5B summarise the results of the mediation analysis. Fornix mediator variables (ISOSF, MPF, *k*_*f*_, R_1_) in Model B fully mediated the age effects on right hippocampal ISOSF, left hippocampal *k*_*f*_ and left hippocampal R_1_ (highlighted in red) but hippocampal mediators in Model A, although demonstrating significant indirect effects on the direct effects of age on fornix R_1_ and *k*_*f*_, did not remove the age effects on fornix MPF, R_1_ and *k*_*f*_. A bi-directional relationship was observed between right hippocampal ISOSF and fornix ISOSF, with both variables fully mediating each other’s age effect. Finally, fornix ISOSF contributed but did not fully mediate the correlation between age and left hippocampal ISOSF and left hippocampal ISOSF did not contribute significantly to any age correlations in fornix metrics. To summarise, full mediation of age effects on glia metrics was observed in Model B but not in Model A (Figure 1), such that age-related glia differences in the fornix mediated age differences in the hippocampus but not *vice versa*.

### Effects of genetic and lifestyle risk factors on age-mediator variables

We explored whether genetic and lifestyle risk factors had an effect on the significant mediator variables (right hippocampal ISOSF, fornix ISOSF, fornix MPF, fornix R_1_ and fornix *k*_*f*_) with hierarchical regression analysis. First age, ICV, sex and years of education were entered into the regression model followed by the stepwise inclusion of all genetic and lifestyle risk variables. Table 2 summarises the results of these regression analyses. Besides age, sex was a significant predictor for differences in fornix and right hippocampal ISOSF. WHR, alcohol consumption and ICV contributed significantly to differences in fornix R_1_. There were also trends (significant on the uncorrected level) for WHR contributions to fornix MPF and *k*_*f*_ and for family history to right hippocampal ISOSF (Table 2).

**Figure 6.**
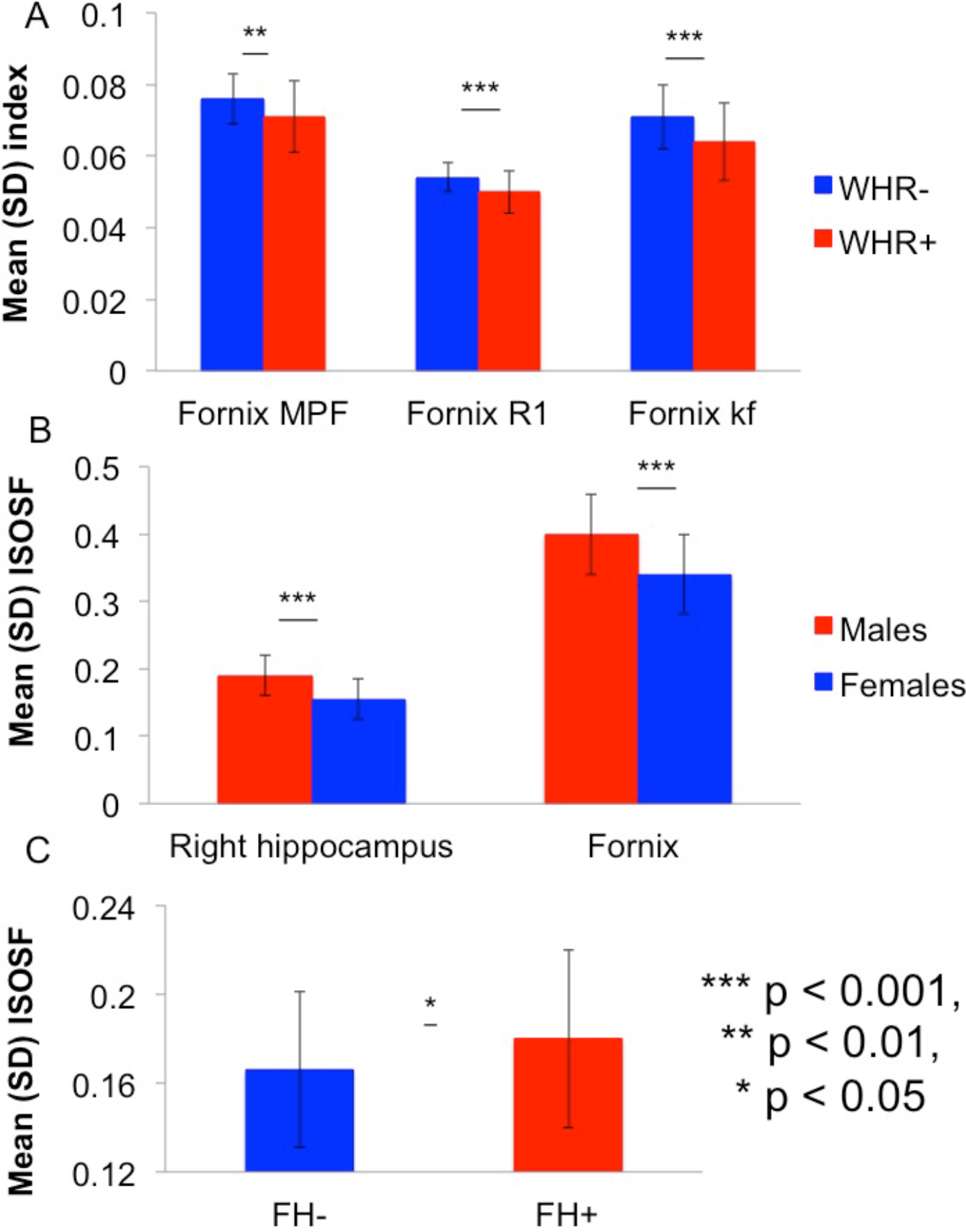
A) Effects of Waist Hip Ratio (WHR) on fornix macromolecular proton fraction (MPF), forward exchange rate *k*_*f*_ and longitudinal relaxation rate R_1_. Centrally obese (WHR+) compared with centrally healthy (WHR−) individuals showed significantly lower MPF, *k*_*f*_ and R_1_ values reflecting reduced apparent myelination in the fornix. B) Men compared with women exhibited significantly larger isotropic signal fraction (ISOSF) in the right hippocampus and the fornix, reflecting more apparent tissue loss in this group. C) Individuals with a positive family history (FH) of dementia (FH+) had larger ISOSF values in the right hippocampus compared with individuals without a FH (FH−). Note that men and women did not differ with regards to their percentage of FH+ individuals (both groups 35.1%).

**Table 2:**
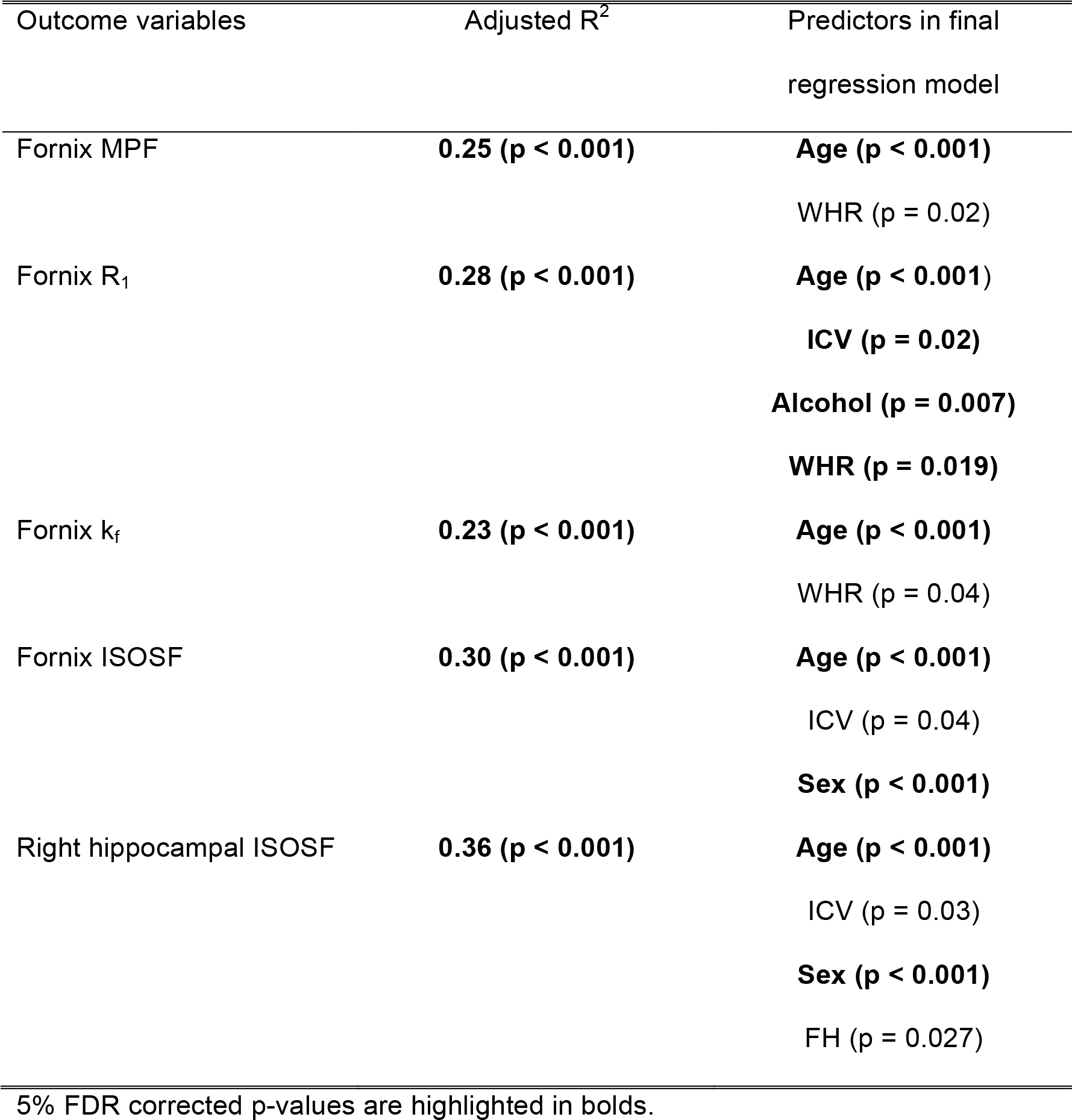
Summary of the results of the hierarchical regression models testing for the effects of genetic and lifestyle risk variables on fornix and hippocampus mediator variables.

*Post-hoc* comparisons revealed that centrally obese individuals compared with individuals with a normal WHR showed lower MPF [t(160) = 2.8, p = 0.005)], R_1_ [t(160) = 3.35, p = 0.0009)] and *k*_*f*_ [t(160) = 4.1, p = 0.00008)] in the fornix. For comparison, there was no effect of WHR on fornix ICSF [t(164) = 1.5, p = 0.13]. Individuals with a positive family history of dementia had higher ISOSF values in the right hippocampus [t(160) = 2.6, p = 0.01)] and men compared with women showed higher ISOSF in the right hippocampus [t(162) = 6.5, p = 0.0000000008)] and in the fornix [t(164) = 6.7, p = 0.00000000002)] (Figure 6). However, there was no difference in fornix R_1_ between individuals that consumed alcohol units above the weekly-recommended limit compared with those within the UK recommended guidelines (p = 0.9).

### Correlations between age-mediator variables and episodic memory performance

Inter-individual differences in delayed verbal recall were negatively correlated with differences in fornix ISOSF [r(164) = −0.22, p = 0.005) and differences in right hippocampal ISOSF [r(164) = - 0.2, p = 0.006].

### CSF partial volume effects

To test whether age effects on fornix and hippocampal qMT metrics were driven by CSF partial volume artefacts, ISOSF metrics were used as mediator variables. Fornix ISOSF contributed significantly to all regression models (p < 0.0001) but did not remove the direct age effect on fornix MPF (t = −3.88, p = 0.0002), R_1_ (t = −3.69, p = 0.0003), and *k*_*f*_ (t = −3.88, p = 0.0002). Left PHC ISOSF contributed to the age effect on right PHC R_1_ (p = 0.0005) but did not remove its age effect (t = −2.36, p = 0.019). No effects of ISOSF were observed on the age effect on left PHC R_2_ (t = −3.57, p = 0.0005). Right but not left hippocampal ISOSF contributed significantly to left hippocampal R_1_ (p = 0.003) and *k*_*f*_ (p = 0.03) and removed the direct age effect on left hippocampal R_1_ (p = 0.05) but not on left hippocampal *k*_*f*_ (p = 0.0013).

## Discussion

The main aim of this study was to discriminate between two classes of causal models of aging with mediation analysis. The neurodegenerative model predicted that age-related hippocampal differences would account for age effects on fornix metrics whilst the neuroglia model predicted that age differences in fornix glia would cause age-differences in the hippocampus. Our results unambiguously support the neuroglia model as we found that fornix glia sensitive metrics of MPF, R_1_ and *k*_*f*_ fully mediated the effects of age on hippocampal tissue properties (Figure 5B) but not *vice versa,* i.e., hippocampal mediator variables, although demonstrating significant indirect effects, did not remove the age effects on fornix MPF, R_1_ and *k*_*f*_ (Figure 5A). This pattern of results is consistent with a growing body of evidence pointing towards an important role of neuroglia changes in aging and neurodegeneration ^14, 16–18, 66–68^. However, to the best of our knowledge, this is the first study to show that age-dependent fornix glia damage mediates hippocampal tissue decline in the human brain.

We observed age-related reductions in fornix MPF, R_1_ and *k*_*f*_, right PHC R_1_ and left PHC R_2_. These metrics are known to be sensitive to myelin, neuroinflammation and to a lesser degree to iron changes in white matter ^32–35, 43, 49, 69, 70^. These effects were not fully explained by ISOSF, thus, do not simply reflect age-related increases in CSF partial volume contamination^71^. In stark contrast, no age effect was present for ICSF from the NODDI model, suggesting that aging did not affect apparent axon density, i.e. the number and the size of axons. A small positive age effect on fornix ODI, however, suggested that fiber coherence may decline with age.

These results provide novel *in vivo* neuroimaging evidence consistent with neuropathological findings of a reduction of up to 45% of the length of myelinated fibers across the lifespan rather than a loss of axons *per se* ^72–76^. Similarly, age-related reductions in fornix myelination and glia changes in the absence of any changes of the fornix cross-sectional area were also observed in non-human primates ^77^. Myelin damage is closely linked to neuroinflammation and qMT metrics are sensitive to both processes^78^. Thus, the here observed age-related differences in MPF and *k*_*f*_ are also consistent with evidence of age-dependent dystrophic and reactive microglia changes^44, 79^.

Currently, a causal link between age-related neuroglia changes and the development of LOAD pathology in the human brain remains speculative. However, white matter disease, characterised by a loss of myelin, oligodendrocytes, axons and reactive astrocyte gliosis ^80–82^ is a known feature of LOAD. Accumulating evidence also suggests that neuroinflammation and a reduction of glia mediated clearance mechanisms contribute to LOAD^14, 15, 83^. For instance, reactive microglia were found in the hippocampus of LOAD brains ^84^ and microglia derived ASC protein specks have been shown to cross-seed amyloid-β plaques in transgenic double-mutant APPSwePSEN1dE9 mice ^85^. It is therefore possible that age-related neuroglia changes involving myelin and inflammation pathways may not only occur in response to neuropathology but may even trigger protein abnormalities and synaptic and neuronal loss. Thus, an intriguing interpretation of our results is that aging leads to glia changes including loss of myelin in the fornix, which in turn may cause tissue damage in the hippocampus that might predispose this region to the development of LOAD pathology ^84, 86^. Clearly, future longitudinal prospective studies assessing the predictive value of MRI indices of white matter glia for amyloid and tau burden are now required to test this hypothesis.

We also observed a bi-directional mediation effect between right hippocampal ISOSF and fornix ISOSF, such that both variables totally mediated the age effect on each other. This result is unsurprising and reflects that tissue loss in the hippocampus is associated with tissue loss in the fornix and the other way around. Reductions in hippocampal ISOSF are consistent with neuropathological findings of a loss of hippocampal pyramidal cells and dentate gyrus granule cells associated with normal aging ^87^. Age reductions in hippocampal R_1_ were accounted for by increases in hippocampal ISOSF and may, therefore, reflect increases in the water content of hippocampal tissue due to loss of macromolecular, neuronal and synaptic tissue and, to a smaller extent, changes in iron concentration. Thus, age-related differences in hippocampal ISOSF and R_1_ are consistent with both glia and neurodegenerative changes in the hippocampus^47, 88^. Importantly though, age-related differences in fornix and hippocampal ISOSF were negatively associated with individual differences in delayed verbal recall performance, suggesting that tissue differences in these regions have an impact on episodic memory ability in cognitive healthy individuals. Age reductions in hippocampal *k*_*f*_, a marker of the rate of the magnetization transfer between macromolecular and free water pool, were not accounted for by hippocampal ISOSF. Giulietti et al. ^45^ found *k*_*f*_ reductions in the hippocampus, temporal lobe, posterior cingulate and parietal cortex in LOAD and proposed that these changes may reflect reductions in metabolic mitochondrial activity ^89^. As age-related reductions of hippocampal metabolic activity have also been observed in animal studies ^90, 91^, the decline in hippocampal *k*_*f*_ observed here may reflect reduced metabolic activity related to aging. Age-related damage to mitochondria of microglia has also been linked to sustained microglia neuroinflammation ^92^.

The question arises how genetic and lifestyle risk factors of LOAD impact on medial temporal tissue properties. Consistent with our hypothesis, obese individuals exhibited reductions in fornix MPF, R_1_ and *k*_*f*_ (Figure 6A). Obesity-related reductions in MRI metrics of proton density, R_1_, and R_2_^*^, have previously been observed in white matter pathways connecting frontal and limbic regions in young adults ^93^. Obesity has also been associated with accelerated aging of cerebral white matter ^62^ and we previously reported Body Mass Index related increases in mean and axial diffusivities in the fornix ^63^. Although the mechanisms underpinning these changes remain unclear, it seems plausible that the observed MRI differences may reflect changes in neuroglia that may be linked with obesity-related systemic inflammation. Consistent with this interpretation, differences in white matter fractional anisotropy in obesity were reported to be associated with inflammation and to a lesser degree with glucose regulation, whilst lower blood pressure and dyslipidemia appeared to have positive effects on white matter microstructure ^94^.

Unexpectedly and in contrast to previous reports ^58, 59, 95^ we did not observe a main effect of the *APOE* genotype on hippocampal gray matter or fornix/PHC white matter tissue properties. There is substantial evidence that *APOE* ε4 carrier status in older individuals (>65 years of age) is related to accelerated atrophy in the hippocampus, increased beta-amyloid burden and to an earlier onset of dementia ^96, 97^. However, the effect size of *APOE*-ε4 genotype on age-related hippocampal atrophy over the adult life course is small (e.g. β= −0.05, p = 0.04 in n = 3749 between 43-69 years of age)^98, 56, 99–102^ and the biological mechanisms underpinning these relationships remain poorly understood. The *APOE* gene is involved in many complex functions, including lipid and amyloid-β metabolism, neuroinflammation and vascular regulation ^103^. It is likely that *APOE* genotype, rather than exhibiting straightforward main effects, interacts with age and with multiple genetic and lifestyle-related factors. As the effects of *APOE* genotype were not the main focus of this study, testing for potential interaction effects between *APOE* genotype and other demographic and risk factors was beyond the scope of this paper.

In contrast to *APOE*, we observed, however, a main effect of family history of dementia on the hippocampus as individuals with a positive history exhibited larger ISOSF in the right hippocampus than those with a negative history (Figure 6C). We also found that males compared with females showed larger ISOSF in the right hippocampus and in the fornix (Figure 6B). These results are compatible with previous findings, indicating that both family history ^104^ and being male ^105–108^ are associated with accelerated age-related atrophy of cortical and subcortical brain regions, including the hippocampus and the fornix.

In conclusion, we provide novel evidence in support of the neuroglia model of aging and LOAD. We propose that age-related damage to fornix glia, rather than a loss of axons, cause hippocampal tissue changes associated with reduced metabolism, glia and neuronal loss. Whilst central obesity also contributed to fornix glia damage, having a positive family history and being male appear to adversely affect hippocampal gray matter. These results suggest that MRI metrics sensitive to neuroglia may potentially provide useful biomarkers of midlife risk of LOAD and, importantly, that healthy lifestyle interventions aimed to reduce the risk of glia damage may reduce the risk of LOAD.

## Methods

This study was approved by the School of Psychology Research Ethics Committee at Cardiff University (EC.14.09.09.3843R2) and all participants gave written informed consent in accordance with the Declaration of Helsinki.

### Participants

Participants were recruited for the Cardiff Aging and Risk of Dementia Study (CARDS) from the Psychology community panel and the employee notice board at Cardiff University and *via* internet and poster advertisements from the local community. Participants had to be community-dwelling individuals between 35 and 75 years of age with a good command of the English language and without a history of neurological disease (e.g. Multiple Sclerosis, Parkinson’s disease, Huntington’s disease), psychiatric disease (e.g. schizophrenia, bipolar disorder, depression requiring hospitalization or a current PHQ-9 score of > 15 indicating severe depression), moderate to severe head injury with loss of consciousness, drug or alcohol dependency, high risk cardio-embolic source (mitral or severe aortic stenosis, severe heart failure, cardiac aneurysm), known significant large-vessel disease (i.e. more than 50% stenosis of carotid or vertebral artery, known peripheral vascular disease, coronary bypass or angioplasty) and MRI contraindications (e.g. pacemaker, cochlear implants, metal pins, stents, screws and/or claustrophobia).

Demographic and health information including information about genetic and lifestyle risk factors of dementia were collected for 211 volunteers of which n = 166 underwent MRI scanning and cognitive testing at the Cardiff University Brain Research Imaging Centre (CUBRIC). Table 1 provides a summary of the demographic, cognitive, and genetic and lifestyle risk information available for these 166 participants.

### Assessment of genetic and lifestyle related risk factors

#### *APOE* genotyping

Participants provided a saliva sample using the self-collection kit “Oragene-DNA (OG-500) (Genotek) for DNA extraction and APOE genotyping. APOE genotypes ε2, ε3 and ε4 were determined by TaqMan genotyping of single nucleotide polymorphism (SNP) rs7412 and KASP genotyping of SNP rs429358. Genotyping was successful in a total of 207 participants including 165 out of the 166 individuals that had undergone an MRI scan. The genotypic distribution of those successfully genotyped can be found in Table 1 and is comparable to the expected frequencies in the normal population ^109^. In addition, participants provided information about their family history (FH) of dementia, i.e. whether a first-grade relative (parent or sibling) was affected by LOAD, vascular dementia or Lewy body disease with dementia.

The following lifestyle related risk factors were documented (Table 1): Abdominal adiposity was assessed by measuring participants’ waist and hip circumferences to calculate the waist-hip-ratio (WHR) following the World Health Organisation’s recommended protocol ^110^. Abdominal obesity was defined as a WHR ≥ 0.9 for males and ≥ 0.85 for females. Systolic and diastolic blood pressure (BP) was measured with a digital blood pressure monitor (Model UA-631; A&D Medical, Tokyo, Japan) whilst participants were comfortably seated with their arm supported on a pillow. The average of three BP readings was taken and hypertension was defined as systolic BP ≥ 140 mm Hg. Other cardio-vascular risk factors of diabetes mellitus, high levels of blood cholesterol controlled with statin medication, history of smoking and weekly alcohol intake in units were self-reported by participants in a medical history questionnaire ^63^. Information about participants’ physical activity over the last week was collected with the short version of the International Physical Activity Questionnaire (IPAQ) ^111^. The median number of hours of non-sedentary activities including walking, gardening, housework and moderate to vigorous activities were recorded. Participants’ verbal intellectual function were assessed with the National

Adult Reading Test (NART) ^112^, immediate and delayed verbal recall abilities with the Rey Auditory Verbal Learning test (RAVLT) ^64^, immediate and delayed non-verbal-recall performance with the Rey Complex Figure^65^, cognitive impairment was screened for with the Mini Mental State Exam (MMSE) ^113^ and depression with the Patient Health Questionnaire for Depression (PHQ-9) ^114^. All participants were cognitively healthy and scored at superior level of intelligence in the NART. Eight participants scored ≥ 10 in the PHQ-9 suggesting moderate levels of depression but no participant was severely depressed.

### MRI data acquisition

MRI data were acquired on a 3T MAGNETOM Prisma clinical scanner (Siemens Healthcare, Erlangen, Germany) equipped with a 32-channels receive-only head coil at CUBRIC.

#### Anatomical MRI

T_1_-weighted anatomical images were acquired with a three-dimension (3D) magnetization-prepared rapid gradient-echo (MP-RAGE) sequence with the following parameters: 256 × 256 acquisition matrix, TR = 2300 ms, TE = 3.06 ms, TI = 850ms, flip angle θ = 9°, 176 slices, 1mm slice thickness, FOV = 256 mm and acquisition time of ~ 6 min.

#### High Angular Resolution Diffusion Imaging (HARDI)

Diffusion data (2 × 2 × 2 mm voxel) were collected with a spin-echo echo-planar dual shell HARDI ^26^ sequence with diffusion encoded along 90 isotropically distributed orientations (30 directions at b-value = 1200 s/mm^2^ and 60 directions at b-value = 2400 s/mm^2^) and six non-diffusion weighted scans with dynamic field correction and the following parameters: TR = 9400ms, TE = 67ms, 80 slices, 2 mm slice thickness, FOV = 256 × 256 × 160 mm, GRAPPA acceleration factor = 2 and acquisition time of ~15 min.

#### Quantitative magnetization transfer weighted imaging (qMT)

An optimized 3D MT-weighted gradient recalled-echo sequence ^115^ was used to obtain magnetization transfer-weighted data with the following parameters: TR = 32 ms, TE = 2.46 ms; Gaussian MT pulses, duration t = 12.8 ms; FA = 5°; FOV = 24 cm, 2.5 × 2.5 × 2.5 mm^3^ resolution. The following off-resonance irradiation frequencies (Θ) and their corresponding saturation pulse amplitude (ΔSAT) for the 11 MT-weighted images were optimized using Cramer-Rao lower bound optimization: Θ = [1000 Hz, 1000 Hz, 2750 Hz, 2768 Hz, 2790 Hz, 2890 Hz, 1000 Hz, 1000 Hz, 12060 Hz, 47180 Hz, 56360 Hz] and their corresponding ΔSAT = [332°, 333°, 628°, 628°, 628°, 628°, 628°, 628°, 628°, 628°, 332°]. The longitudinal relaxation time, T_1_, of the system was estimated by acquiring a 3D gradient recalled echo sequence (GRE) volume with three different flip angles (θ = 3,7,15). Data for computing the static magnetic field (B_0_) were collected using two 3D GRE volumes with different echo-times (TE = 4.92 ms and 7.38 ms respectively; TR= 330ms; FOV= 240 mm; slice thickness 2.5 mm) ^116^.

T_2_-weighted maps were acquired with a multi-echo spin echo sequence with five equally spaced echo times (TEs = 13.8ms – 69ms) and the following parameters: TR = 1600ms, 19 slices, 5 mm slice thickness, FOV = 240mm, flip angle θ = 180° and acquisition time of ~ 6 min.

### MRI data processing

The two-shell diffusion-weighted HARDI data were split and b = 1200 and 2400 s/mm^2^ data were corrected separately for distortions induced by the diffusion-weighted gradients and artifacts due to head motion with appropriate reorientation of the encoding vectors ^117^ in ExploreDTI (Version 4.8.3) ^50^. EPI-induced geometrical distortions were corrected by warping the diffusion-weighted image volumes to the T_1_–weighted anatomical images which were down-sampled to a resolution of 1.5 × 1.5 ×1.5 mm ^118^. After preprocessing, the Neurite Orientation Dispersion and Density (NODDI) model ^31^ was fitted to the dual-shell HARDI data using fast, linear model fitting algorithms of the Accelerated Microstructure Imaging via Convex Optimization (AMICO) framework ^119^ to obtain isotropic signal fraction (ISOSF), intracellular signal fraction (ICSF) and orientation dispersion index (ODI) maps (Figure 2).

MT-weighted GRE volumes for each participant were co-registered to the MT-volume with the most contrast using a rigid body (6 degrees of freedom) registration to correct for inter-scan motion using Elastix ^120^. The 11 MT-weighted GRE images and T1-maps were modelled by the two pool Ramani’s pulsed MT approximation ^121^. This approximation provided maps of the macromolecular proton fraction (MPF), the forward exchange rate *k*_*f*_ and the spin lattice relaxation rate of the free pool *R*_1_. MPF maps were threshholded to an upper intensity limit of 0.3 and *k*_*f*_ maps to an upper limit of 3 using the FMRIB’s fslmaths imaging calculator to remove voxels with noise-only data.

A monoexponential decay function was fitted to the T_2_ images using the quimultiecho program from the Quantitative Imaging Tools (QUIT) library (http://spinicist.github.io/QUIT). This fitting excluded the first acquired TE due to a non-contained stimulated echo artifact in the first echo of the multi-contrast spin echo Siemens library sequence. R_2_ was calculated as 1/T_2_ in second units.

All image modality maps and region of interest masks were spatially aligned to the T_1_-weighted anatomical volume as reference image with linear affine registration (12 degrees of freedom) using FMRIB’s Linear Image Registration Tool (FLIRT).

### Tractography

The RESDORE algorithm ^122^ was applied to identify outliers, followed by whole brain tractography with the damped Richardson-Lucy algorithm (dRL) ^123^ on the 60 direction, b = 2400 s/mm^2^ HARDI data for each dataset in single-subject native space using in house software ^122^ coded in MATLAB (the MathWorks, Natick, MA). To reconstruct fibre tracts, dRL fibre orientation density functions (fODFs) were estimated at the center of each image voxel. Seed points were positioned at the vertices of a 2×2×2 mm grid superimposed over the image. The dRL tracking algorithm interpolated local fODF estimates at each seed point and then propagated 0.5mm along orientations of each fODF lobe above a threshold peak of 0.05. This procedure allowed four potential streamlines to emanate from each seed point. Individual streamlines were subsequently propagated by interpolating the fODF at their new location and propagating 0.5mm along the minimally subtending fODF peak. This process was repeated until the minimally subtending peak magnitude fell below 0.05 or the change of direction between successive 0.5mm steps exceeded an angle of 45°. Tracking was then repeated in the opposite direction from the initial seed point. Streamlines whose lengths were outside a range of 10mm to 500mm were discarded.

The fornix and the parahippocampal cinguli were reconstructed with an in-house automated segmentation method based on principal component analysis (PCA) of streamline shape ^124^. This procedure involves the manual reconstruction of a set of tracts that are then used to train a PCA model of candidate streamline shape and location. Twenty datasets were randomly selected as training data. The fornix and the PHC tracts were reconstructed by manually applying waypoint region of interest (ROI) gates (“AND”, “OR” and “NOT” gates following Boolean logic) to isolate specific tracts from the whole brain tractography data. ROIs were placed in HARDI data native space on colour-code fiber orientation maps in ExploreDTI following previously published protocols ^24, 63, 125, 126^. The trained PCA shape models were then applied to all datasets: candidate streamlines were selected from the whole volume tractography as those bridging the gap between estimated end points of the fornix or the PHC and spurious streamlines were excluded by means of a shape comparison with the trained PCA model. All automatic tract reconstructions underwent quality control through visual inspection and any remaining spurious fibers that were not consistent with the tract anatomy were removed from the reconstruction where necessary.

### Whole Hippocampal segmentation

Volumetric segmentation of the left and right whole hippocampi from T_1_-weighted images was performed with the Freesurfer image analysis suite (version 5.3), which is documented online (https://surfer.nmr.mgh.harvard.edu/). Mean intracranial volume fractions (ICV) were extracted for each brain as estimates of individual differences in headsize.

### Statistical analyses

Statistical analyses were conducted in SPSS version 20 ^127^ and the PROCESS computational tool for mediation analysis ^128^. Multiple comparisons were corrected with a False Discovery Rate (FDR) of 5% using the Benjamini-Hochberg procedure^129^. Partial Eta^2^ (ηp^2^) and correlation coefficients are reported as indices of effect sizes. Differences in the size of correlation coefficients were tested with the Fisher’s r-to-z transformation ^130^. All reported p-values are two-tailed.

Omnibus multivariate regression analysis was conducted to test for main effects of age, genetic (*APOE* genotype and family history of dementia) and lifestyle related risk factors (WHR, systolic and diastolic blood pressure, weekly alcohol consumption and physical activity) (Table 1) as well as potentially confounding variables of sex, education and head size simultaneously on all white and gray matter MRI outcome metrics (Figure 2). Given the low numbers of diabetics, smokers, and individuals on statins amongst our sample (Table 1), these variables were not included in the statistical analyses. Omnibus effects were followed up with *post-hoc* multivariate covariance analysis testing for effects on individual MRI metrics in the five regions of interest (left and right hippocampi, left and right parahippocampal cinguli, fornix) while correcting for any confounding variables.

The direction of age effects and the relationship between hippocampal and white matter microstructural metrics were further investigated with partial Spearman correlation coefficients controlling for confounding variables (sex and ICV). Non-parametric Spearman correlation was chosen as sex is a categorical variable and the calculation of parametric Pearson correlations requires continuous variables.

Mediation analysis was then carried out to test for the indirect effects a*b of hippocampal mediator variables on the direct effects c’ of age on the fornix metrics (Figure 5 Model A) and for indirect effects a*b of fornix mediator variables on direct age effects c’ on hippocampal differences (Figure 5 Model B). The significance of indirect and direct effects was assessed with a 95% confidence interval based on bootstrapping with 5000 replacements (89).

The impact of genetic and lifestyle risk factors on individual mediator variables, was then explored with hierarchical linear regression analyses. These included in a first model age and all other potentially confounding variables and in a second model added all genetic and lifestyle risk factors in a stepwise fashion to the model. Group differences in hippocampal and fornix mediator variables for significant predictor variables were assessed with independent t-tests. Finally, Pearson correlation coefficients between MRI mediator variables and immediate and delayed, verbal and non-verbal recall performance were calculated to explore brain-function relationships.

## Acknowledgements

This research was funded by a Research Fellowship awarded to Claudia Metzler-Baddeley from the Alzheimer’s Society and the BRACE Alzheimer’s Charity (grant ref: 208). DKJ is supported by a Wellcome Trust Investigator Award (096646/Z/11/Z) and a Wellcome Trust Strategic Award (104943/Z/14/Z). We would like to thank Erika Leonaviciute, Peter Hobden and Sonya Foley-Bozorgzad for their assistance with MRI data acquisition and processing and Rosie Dwyer, Samantha Collins, Abbie Stark, and Emma Blenkinsop for their assistance with the collection and scoring of the cognitive and health data.

We also would like to thank Rhodri Thomas for his assistance with the *APOE* genotyping of the saliva samples.

The authors declare no competing financial or non-financial interests.

## Author contributions

CM-B is the PI of the study and is responsible for the conceptualization and data acquisition and analyses of the study. CM-B has also written the manuscript. JPM was responsible for participant recruitment, data acquisition and MRI data processing. RS was responsible for the *APOE* genotyping. FF and JE have prepared the qMT and diffusion MRI protocols and have helped with MRI data processing. RJB was involved in the conceptualisation and has advised on statistical data analysis. JPA and DKJ and all other authors have reviewed the manuscript.

